# SODAR: managing multi-omics study data and metadata

**DOI:** 10.1101/2022.08.19.504516

**Authors:** Mikko Nieminen, Oliver Stolpe, Mathias Kuhring, January Weiner, Patrick Pett, Dieter Beule, Manuel Holtgrewe

## Abstract

Scientists employing omics in life science studies face challenges such as the modeling of multi assay studies, recording of all relevant parameters, and managing many samples with their metadata. They must manage many large files that are the results of the assays or subsequent computation. Users with diverse backgrounds, ranging from computational scientists to wet-lab scientists, have dissimilar needs when it comes to data access, with programmatic interfaces being favored by the former and graphical ones by the latter.

We introduce SODAR, the system for omics data access and retrieval. SODAR is a software package that addresses these challenges by providing a web-based graphical user interface for managing multi assay studies and describing them using the ISA (Investigation, Study, Assay) data model and the ISA-Tab file format. Data storage is handled using the iRODS data management system, which handles large quantities of files and substantial amounts of data. SODAR also offers programmable APIs and command line access for metadata and file storage.

SODAR supports complex omics integration studies and can be easily installed. The software is written in Python 3 and freely available at https://github.com/bihealth/sodar-server under the MIT license.

## 1. Introduction

Modern studies in life sciences rely on “omics” assays, which encompass branches of science such as genomics, proteomics, and metabolomics. One or multiple assays can be run within a single study, potentially including assays for multiple omics studies of several types.

The following key steps are required for executing these complex omics studies: a) planning which results in study metadata; b) collection of mass data; and c) data analysis, including the integration of multiple assays. The aim of SODAR is to ensure support for scientists within all the steps.

### 1.1 Challenges

Each step presents its own set of challenges. During planning it is important to enable recording crucial factors and covariates. The flow of materials and samples through processes must also be specified in sufficient detail. Further challenges arise from, e.g., assays using complex multiplexing, such as the need for reference samples; requirements for using controlled vocabularies or ontologies; and possible change of assays over time.

In the data collection step, scientists must record the used machines, kits, and versions of both hardware and software used. Omics studies also create large volumes of data, ranging from a few gigabytes for mass spectrometry to terabytes for imaging such as microscopy. This data may be spread among many files, further complicating the needs for managing mass data storage. Instead of a rigid process, data collection should also be adjustable to changes and developments in data generation over time.

Data analysis is often split into multiple phases, with primary analysis of each assay followed by steps for integration of results. Specific results need to be fed back to metadata management, annotation, quality control or storing resulting markers. Access to metadata with recorded factors and confounders is necessary in each step, while access to primary raw data becomes less important after the primary analysis. Certain analysis results are written back into the mass data storage. This includes binary alignment map (BAM) files, and variant call format (VCF) files.

There are also overarching challenges for the steps in study execution. All data should be recorded in structured format. Automation should be applied where possible, and on-premise installation might be preferable or even required when data privacy relevant data is generated such as DNA sequencing.

### 1.2 Data Management Approaches

In this and the following section, we will discuss the topic of data management and software. The terms “data” and “document” will be used interchangeably in this section. The steps described in section 1.1 can be interpreted as processes taking documents and materials as input, and generating more documents and materials as the result. For example, data collection takes the plan document and samples and generates assay result files (documents). Scientists thus need computational tools for supporting them in managing their scientific and research data.

Historically, such documents are maintained on paper in laboratory notebooks, or documentation created by quality control systems. For the most direct and unstructured approaches in maintaining digital data, this corresponds to word processing, spreadsheet, and image files on local or network drives. More structured approaches are desireable for taking advantage of digital documents, preventing research data loss [1] or fostering re-use [2].

While data management in science is a broad topic, the library and information science community is frequently approaching it using a top-down approach. Frequently, in this context, the term “research data management” (RDM) is used. Here, the needs of whole organizations and their parts for managing their resarch data, as well as the necessary steps to establish whole RDM systems are considered first, for example cf. Donner [3]. This correlates with the role of libraries in certain academic organizations for organizing data that was collected in research.

A second approach which can be described as “bottom-up” originates from different “working scientist” communities. The communities commonly refer to the topic as “scientific data management” (SDM) and solve their problems at hand, often starting with specific small-scale solutions which are then upscaled if the need arises. While considering their organizational embedding, they focus on solving specific data management challenges for themselves and their peers. We found ourselves in this situation and will thus focus on this perspective.

### 1.3 Data Management Software Packages

Scientific data management needs come in different forms and shapes. We could find no general treatment of the subject of data management in the literature. Machina and Wild [4] provide a collection of four tool categories: laboratory information management systems (LIMS), electronic laboratory notebooks (ELN), scientific data management systems (SDMS) and a chromatography data system that we generalize as instrument-specific data system (IDS). In this section, we provide our take on explaining what these systems comprise. We also note – as Machina and Wild [4] did – that categorization of such software solutions is not clear-cut, and features may be overlapping. We expand this list by two more system types: data repository systems (DRS) and database/data warehouse management frameworks (DMF).

The four items by Machina and Wild [4] are as follows:

**LIMSs** focus on storing information around laboratory workflows. This includes tracking of consumables, samples, instruments, and tests. They deal with daily tasks of laboratories such as billing and instrument calibration. They are often specific to certain domain areas such as sequencing facilities.

**ELNs** focus on allowing humans to record their laboratory work. They replace paper notebooks and capture experiments and their results, mostly in free-form text, pictures, tables etc. They play a key role in fulfilling regulatory requirements.

**IDSs** provide data capturing, storage, and analysis functionality in instrument-specific domains. Two examples are the CASAVA pipeline and the BaseSpace cloud-based service, both from Illumina. The former is provided without extra cost with the instrument along with its source code, while the latter is purchasable and closed source. Such software often ships with the instruments themselves.

**SDMSs** provide scientific content management functionality for scientific data and documentation. They allow for the management of metadata and potentially mass data. Their core functionality does not include data analysis, user-centric data collection, or laboratory workflow tracking. Such features may be potentially supported by plugins or extensions. Many such systems offer integration with surrounding systems, e.g., via application programming interfaces (APIs).

We augment this list by two system types:

**DRSs** provide shared access to data with appropriate documentation and metadata. Examples are FAIRdom Seek [5], Dataverse [6], and Yoda [7]. There also specialized DRS focusing on particular use cases such as dbGAP [8], MetaboLights [9], and Gene Expression Omnibus [10], that allow for managing public or controlled public access to large research data collections.

**DMFs** allow for the rapid development of database and data warehouse applications. They often provide pre-existing components to build on readymade functionality and extension by implementing custom components. Such enable creating domain-specific databases and structured data capturing. Examples include Molgenis [11] and Zendro [12].

Other types of systems also exist and not every system falls into just one category. A complete review of such systems is beyond the scope of this manuscript. This section identifies focus areas of systems involved in some form of scientific data management. SODAR falls into the category of SDMS.

### 1.4 Data Management Technologies

For planning and documenting experiments and their structure, experiment oriented metadata storage formats with predefined syntax and semantics exist. A popular standard is the ISA (Investigation, Study, Assay) model [13], which allows describing studies with multiple samples and assays. The ISA model defines the ISA-Tab tabular file format, which allows users to model each processing step with each intermediate result and annotate each of these with arbitrary metadata. An example of an alternative to ISA-Tab is Portable Encapsulated Projects (PEP) [14]. There are also more specialized standards such as Brain Imaging Data Structure (BIDS) for brain imaging data [15], as well as other approaches such as Clinical Data Interchange Standards Consortium (CDISC) standards [16], and the Hierarchical Data Format (HDF5) [17]. Use of generic file formats such as HDF5, TSV, XML and JSON is also common.

For storing large volumes of omics data, it is possible to simply use file systems or object storage systems. More advanced solutions such as Shock [18] or dCache [19] allow for storing metadata and distributing data over multiple servers. iRODS (Integrated Rule-Oriented Data System) [20, 21] adds further features, such as running rules and programs within the data system and enabling integration with arbitrary authentication methods.

For publication, raw and processed data and metadata are deposited in scientific catalogues, study databases and registries. Examples include the BioSamples database for metadata [22] and Sequence Read Archive (SRA) for raw sequencing data [23].

### 1.5 Our Work

In our work, we focus on managing many omics projects of varying data size and various use cases including cancer and functional genomics studies. We also need to support multiple technologies such as whole genome sequencing, single cell sequencing, proteomics, and mass spectrometry. Deposition to public repositories was not a necessity in our context. However, SODAR is an ISA compliant system. Should the data owner require it, it is easily feasible to create appropriate exports to public data repositories using the APIs provided by SODAR. Open source software is a requirement to avoid vendor lock-in and allow for flexibility in different use cases. A suitable end-to-end solution was not available when we started our work in 2016. Therefore, we set out to implement an integrated system for managing omics-specific data and metadata.

In this manuscript, we introduce SODAR (the System for Omics Data Access and Retrieval). SODAR combines the modeling of studies and assays using the ISA-Tab format with handling of mass data storage using iRODS. More example projects are available in the SODAR online demo at https://sodar-demo.cubi.bihealth.org.

## 2. Results

We present the results by first giving an overview of the developed SODAR system. Next, we compare it to a selection of existing tools and their relevant features. We then describe processes we have established around SODAR. Finally, internal usage statistics are detailed along with discussion on the limitations of SODAR.

### 2.1 Resulting System Overview

Figure 1 presents the components of the SODAR system. The SODAR server is built on the Django web framework. It contains the main system logic and provides both a graphical user interface (GUI) and application programming interfaces (API) for managing projects, studies, and data.

Project and study metadata are stored in a PostgreSQL database. The study metadata is stored as ISA-Tab compatible sample sheets, with each project containing a single ISA-Tab investigation. Each investigation can hold multiple studies, likewise each study can contain multiple assays.

Mass data storage is implemented using iRODS and accessed via iRODS command line tools or access to the WebDAV protocol, which is provided by using the Davrods software. The SODAR server manages creation of expected iRODS collections (i.e., directories), governs file access and enforces rules for file uploads and consistency. Investigations, studies, and assays correspond to collections in the iRODS file hierarchy. Within assays, collection structure can be split by, e.g., samples or libraries, depending on the type of assay.

Uploading files for studies is handled using “landing zones”, which are user specific collections with read and write access. The SODAR server handles validation and transfer of files from the landing zones into the project specific read-only sample repository, which is split into assay specific iRODS collections.

Planning and tracking the study design and experiments is done using the ISA-Tab compatible sample sheets. Here, the “assay” in the ISA model corresponds to an “experiment” in our work. SODAR provides multiple ways to create and edit both the metadata model and the contained metadata itself, including user friendly GUI-based creation of sample sheets from ISA-Tab templates. The templates aid in maintaining consistent metadata structures between studies. Once created, the SODAR server provides a graphical UI for filling up metadata and configuring expected values, including support for controlled vocabularies and ontologies. Furthermore, SODAR also allows uploading and updating sample sheets using its API. Uploading any valid ISA-Tab file and replacing existing sheets via upload is also supported, enabling the creation of sample sheets using other software such as ISA-tools [13]. The API allows to automate metadata and file management activities using scripts.

**Figure 1.**
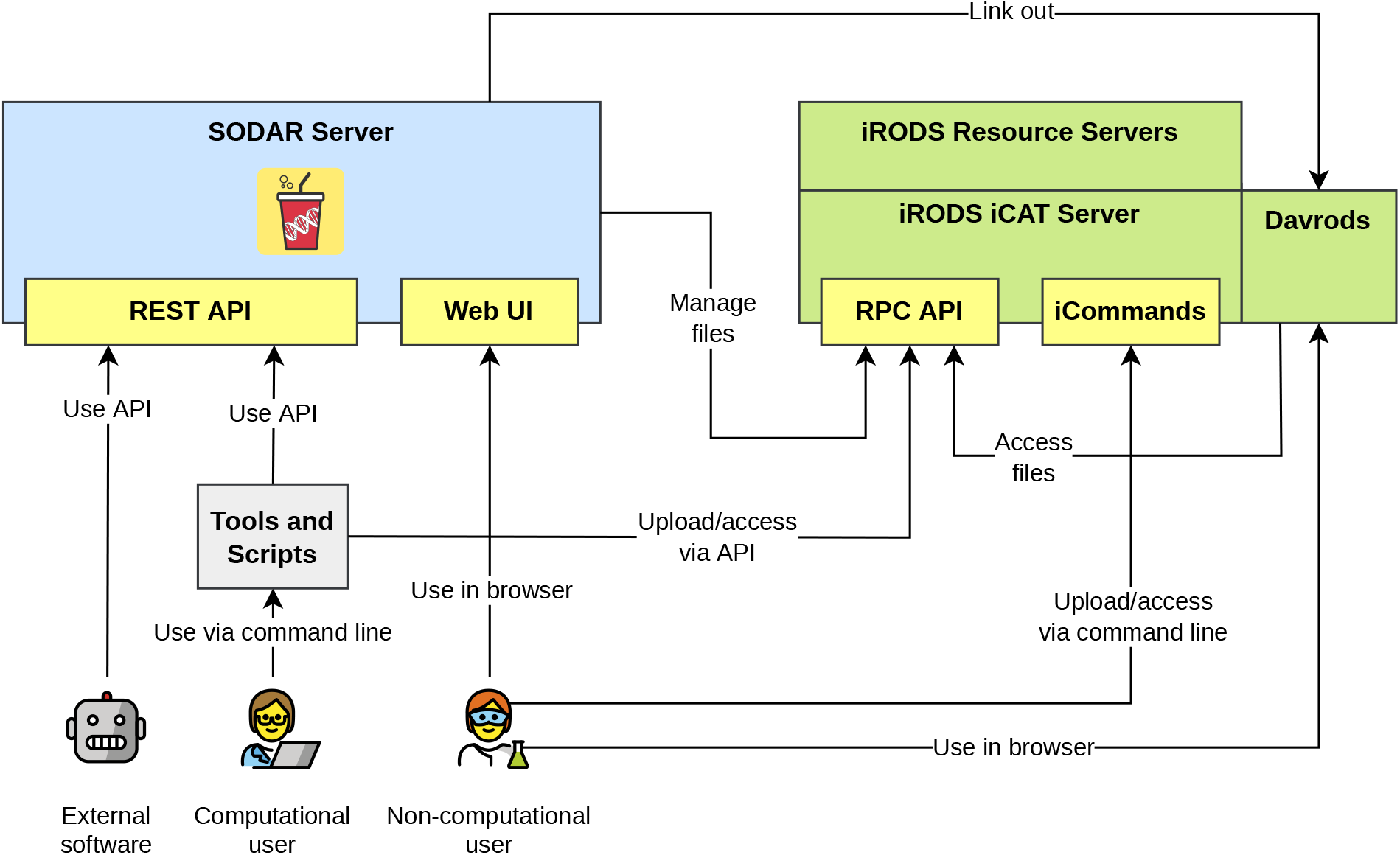
SODAR system with its components and actors. The figure illustrates how actors interact with SODAR and iRODS through different APIs.

### 2.2 Data Management Software Features and Selection

This section first describes features of DMS packages that are subsequently used for comparing SODAR to other software types and packages. We then describe the selection process for software comparison.

The following is a list of features that allows us to see the unique strengths and properties of SODAR in the category SDMS and describe the difference to other categories. When a feature is important in multiple categories, it is only shown once. Categories 1-4 are focused on SDMS, and category 5 contains features also important for other categories.

1. Features addressing overarching challenges

a. Structure into projects and folders
b. Access control
c. Automation possible via API
2. Use of open formats and standards

a. Features addressing planning challenges
b. Structured recording of assays and experiments
c. Flexibility in definition of studies and experiments
d. Annotation with controlled vocabulary
e. Annotation with ontologies
3. Features addressing data collection challenges

a. Storage of files possible
b. Support for many files
c. Support for large file sizes
4. Features addressing data analysis challenges

a. API for reading and updating experiment metadata
b. API for reading and updating mass data
5. Features commonly found in specific systems

a. ELN

i. Flexible data entry in free text / tables / pictures
b. DRS

i. Host public data repositories
c. DMF

i. Easy creation of new data tables
ii. User-centric data entry
iii. Multiple predefined components, e.g., for data visualization and analysis

With the aim of showing the unique strengths of software categories and packages, we attempted to select popular software packages in each category. We limited the selection to open-source software. We searched for the different software types via a publication on Google Scholar or the project search on GitHub. We made no attempt to define “the most popular” or “the best” software packages. We excluded LIMS and IDS as such software is focused on the wet-lab process. The following software was selected:

1. SDMS

a. SODAR
b. qPortal [24]
c. FAIRDom Seek [5]
d. OpenBIS ELN-LIMS [25]
2. ELN

a. ELabFTW [26]
3. DRS

a. Dataverse [6]
b. Yoda [7]
4. DMF

a. Molgenis [11]
b. Zendro [12]

### 2.3 Data Management Software Comparison

The table included in [Additional File 1] shows the comparison of the categorized software in the categories as described in section 2.2.

Since the software packages operate in a similar space, there is a certain overlap in features, even across categories. Most software packages provide the features for addressing the overarching challenges. All “planning” features are included in SODAR and FAIRDom Seek in the SDMS category, while qPortal and OpenBIS remain limited. ELabFTW provides limited functionality for structured recording and does not support controlled vocabularies and ontologies, while DRS systems do not address planning challenges by their design. As expected, such features can be implemented by the DMF packages, but they do not provide the functionality on their own. The “data collection” and “data analysis” features are only comprehensively addressed by SODAR and FAIRDom Seek in the SDMS category, with FAIRDom Seek being limited in storing many and/or large files. ELN software is limited in this capability, while DRS packages provided good support for such features, and the DMF software packages allow for implementing support to varying levels.

As for the specialized features, some functionalities of “foreign” categories are implemented. For example, SODAR has support for user centric data entry, and FAIRDom Seek allows for hosting public data repositories by design. However, each software package shows its strengths by providing the features for the tasks that it was originally designed for. We note that certain packages cover their category more focused or comprehensively than others. For example, in the DMF category, Molgenis has an ecosystem of many predefined components, while Zendro focuses on allowing for the easy creation of tables and creating user centric data entry masks.

### 2.4 Roles and Interaction with SODAR

The general workflow in using SODAR for managing data and metadata is shown in Figure 2. We distinguish between the roles “data steward” and “experimentalist.” It is possible for one person to act in both roles.

Data stewards are responsible for creating the overall structure of the experiment data. They are expected to be experienced with using ISA-Tab files. For example, in our use case, data stewards are bioinformaticians working in the core unit. They are responsible for planning the experiments and modeling them in the ISA-Tab format as sample sheets describing the overall experimental design. Data stewards also maintain a library of sample sheet templates for common use cases. With experienced experimentalists the steward might just create the general structure of the experiment. In some cases, the steward may also pre-create the sample sheet with an initial structure of all planned samples and processes and IDs together with experimentalists.

Experimentalists are primarily responsible for entering the actual data into the system. They are users more concerned with completing the metadata in the sample sheet than in creating its structure. When the full sample sheet is created together with data stewards, experimentalists may only verify the structure against the information of their experiments and fill in some measurements in sample sheet cells, e.g., concentration measurements. More experienced experimentalists will also create new rows in the ISA-Tab tables for samples, related materials, and processes.

**Figure 2.**
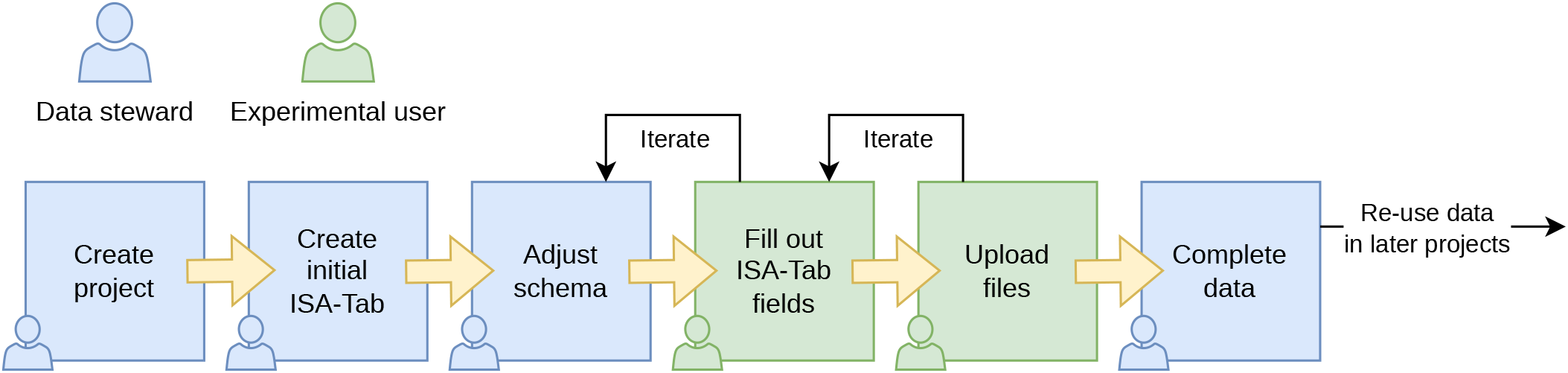
SODAR metadata management workflow. The workflow scheme is divided into steps attributed to a data steward (blue) who manages the overall data schema and experimental user (green) who enters the actual data or uploads files.

### 2.5 General SODAR Process

Here we describe the SODAR-backed process of managing experiment data we are using in our work. This demonstrates how SODAR helps tackle challenges in complex omics study management.

#### 2.5.1 Planning and Sample Sheet Creation

Planning begins with data steward and experimentalists meeting and discussing the study, including, e.g., its factors, sample size, replicas, and confounders. Stewards create sample sheets from templates and modify columns depending on the discussions and the study’s requirements. Working together, stewards and experimentalists also decide on ontologies and controlled vocabularies to use, data ranges, etc.

The template will be bootstrapped with example samples, or all samples, depending on the study. During this step, the experimentalist receives training in using the SODAR sample sheet editor for filling in cells where necessary. Filling cells can involve, e.g., adding measurements, cancer staging, definition and refinement of phenotypes, adjustment of relationship information.

Automated extraction of measurements from instruments or LIMS and ingesting it using the SODAR API is also possible. for example, an integration with a LIMS system could automatically create samples as they are processed in the wet lab, while measurements could be written to SODAR from the LIMS or from an integration of an ELN system. We are currently working towards this when cooperating with other units.

#### 2.5.2 Data Acquisition and Sample Sheet Update

Experimentalists run their experiments and use SODAR for editing the sample sheets. This includes adding new samples, marking dropouts, or removing them, and adjusting ontologies and terms as needed. SODAR sample sheets are useful as a central storage of metadata, removing the need to, e.g., share spreadsheets via email. Differences between sample sheet versions can also be browsed in the SODAR UI to track changes in the metadata.

In this step, actual data files are uploaded by experimentalists to the project sample repository through landing zones. The iRODS collection structure for each study is maintained by SODAR and based on the study type and names of samples or associated libraries. In most cases, files related to a certain sample and its processing in an assay can be found in the collection named after the related library.

#### 2.5.3 Data Analysis

For data analysis, bioinformaticians access metadata in the sample sheets as well as raw data in iRODS, the latter being linked to former in the SODAR GUI for ease of access. Depending on the phase of study, this may involve, e.g., primary analysis, secondary analysis, and required data integration. Resulting files are uploaded back into iRODS via SODAR for safekeeping and sharing between researchers. Also uploaded are files needed for integrating with third party systems, such as UCSC Genome Browser [27] tracks and files for data exploration tools such as SCelVis [28].

During the analysis, up-to-date experiment structure is maintained in SODAR. It represents a centralized storage and sole source of truth for the internal structure, encompassing factor values, ontologies, and controlled vocabularies. Similarly, it represents an external structure, with samples and materials linked to corresponding iRODS collections.

SODAR also provides integrations to specific third-party software to aid analysis. For germline and cancer DNA sequencing experiments, SODAR supports the IGV Genome Browser [29], by generating session files pointing at relevant variant and read alignment files with a single click.

#### 2.5.4 Long-Term Data Storage and Data Access

After transferring files from landing zones into the project’s sample repository, the data is in general assumed to be permanent and not modifiable or rewritable, with users only having the possibility of request file deletion from project maintainer in case of, e.g., mistakes in uploading. Hence, once the project finishes, the data is considered good for long term archival. SODAR supports setting projects into a read-only “archived” state and provides an API for implementing custom policies for handling archived data. For example, such a policy might consist of adding a cold storage resource such as tape onto which the data could be moved.

In exporting data to public databases, creating a generic exporter cannot be considered feasible due to the metadata model flexibility in SODAR. However, there are export possibilities depending on the type of study. For example, if the project is set up with Gene Expression Omnibus (GEO) [10] compatible metadata, exporting to the GEO database may be trivial depending on the target system APIs. In the future, we intend to create export functionality from SODAR to the emerging German National Research Data Infrastructure (NFDI), the associated German Human Genome-phenome Archive (GHGA) [30], and corresponding metadata models. These will be based on the federated European Genome-phenome Archive (EGA) [31] and should provide a good starting point for many other exporters. NFDI will be our long term and controlled public access backend, while other users and instances might have other backends.

### 2.6 Internal Usage Statistics

We have been using SODAR in our group’s projects for the past four years. Table 2 summarizes data statistics and metadata stored in our internal instance and the diversity of projects. We have thus tested SODAR extensively in a real-world setting and use it daily as our main storage for all our project data and metadata.

**Table 2.**
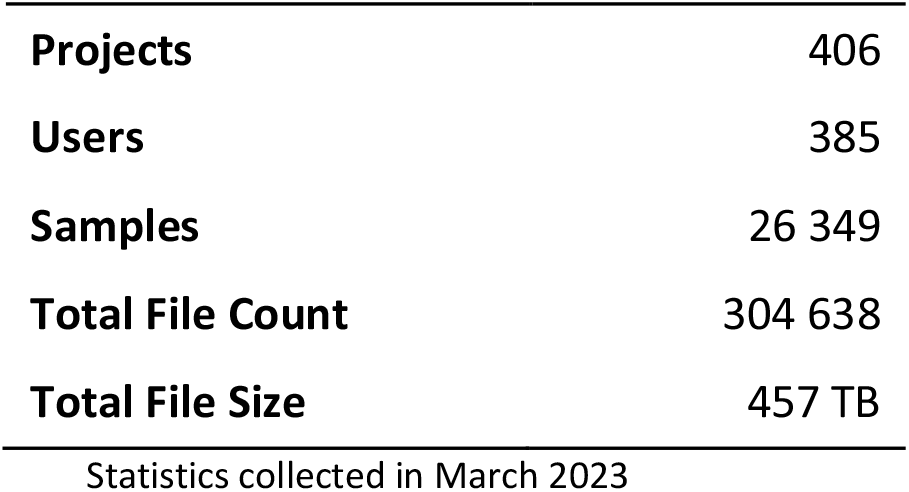
Summary statistics of project type and count, sample count, user count, mass data file count and total size in our internal instance of SODAR.

|  |  |
| --- | --- |
| **Projects** | 406 |
| **Users** | 385 |
| **Samples** | 26 349 |
| **Total File Count** | 304 638 |
| **Total File Size** | 457 TB |

Statistics collected in March 2023

Figure 3 displays file size and count for each project on our system in March 2022. The diagram shows the varying scale of the projects within our group. A limited number of projects between a 20-45 terabyte range can be seen, while most are smaller.

**Figure 3.**
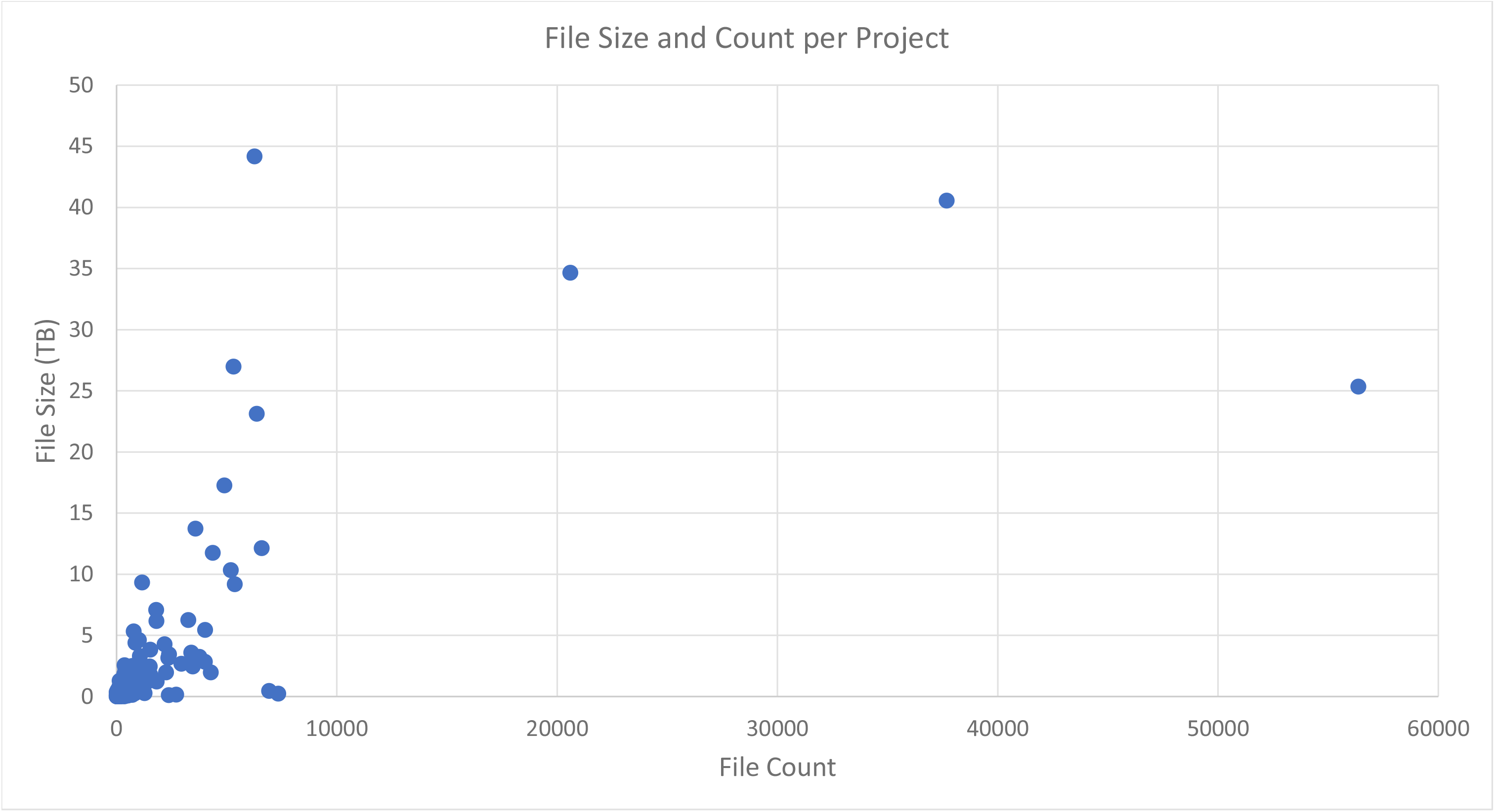
SODAR project file statistics scatter plot, with file count per project on the X axis, and the total file size in terabytes on the Y axis.

### 2.7 Limitations

Currently, SODAR offers no automated data export to, e.g., the GEO database. This may be added in the future as discussed in the “Long-term data storage” section. Similarly, SODAR does not support access in a “data commons” manner. It is possible to set specific projects for public read access, but by default SODAR enforces strict access control to data.

We also do not have a definitive solution for training people in ISA-Tab. SODAR features a set of templates for predefined study types for, e.g., germline and cancer studies, but there is no definite solution for trivially setting up any type of study as ISA-Tab.

## 3. Methods

SODAR is implemented in Python 3 using the Django web framework and Django REST Framework. Reusable components have been extracted into the library SODAR Core [32]. ISA-Tab format manipulation has been implemented using AltamISA [33].

### 3.1 Project Organization, Authorization Structure, and LDAP Integration

SODAR uses the concept of “projects” for organizing all data. Projects have a unique identifier and some basic metadata, such as title and description. Projects are organized in a tree structure using the concept of “categories” that can contain projects or other categories. Each project has a single owner, who can assign themselves a delegate for managing the project. Further users can be granted access to the project either in a read-write (contributor) or a read-only fashion (guest) using Role-Based Access Control (RBAC) [34].

SODAR can be configured to be run standalone or integrated with LDAP servers, including Microsoft ActiveDirectory, for providing authentication information. Here, authentication refers to checking the identity of a user based on their username and password.

### 3.2 iRODS integration

SODAR automatically manages user access to projects in iRODS. This is done by creating an iRODS directory and user group for each project. The group is given access to the directory and group membership is synchronized between the SODAR database and iRODS.

SODAR creates an iRODS collection for each study and assay from the ISA model of the project. Files can be uploaded by users through landing zones, either for each sample or for the whole study or assay. It is thus possible to add data for an arbitrary number of assays for each sample and original donor or specimen.

The files can be accessed either directly through iRODS or using the WebDAV protocol through the Davrods [35] software. The latter allows users to access the storage as a network drive on their desktop computers. Since WebDAV is HTTP based, users can also make data available to genome browsers such as IGV or UCSC Genome Browser. Moreover, it is easy to access data through an organization’s security system and proxies without the intervention of IT departments.

Optionally, SODAR allows the management of iRODS “tickets,” which allow for access based on randomly generated tokens instead of user login. This way, users can upload genome browser tracks to SODAR and iRODS and create public URL strings to access them and share them with users that do not have access to the full project, or do not even have an account in SODAR.

### 3.3 Sample Sheet Editor, Import, Export

Sample sheets can be included into SODAR projects by either importing existing ISA-Tab files or template-based creation. When importing, the user can upload a Zip archive or a set of individual ISA-Tab files. For creating sample sheets from templates, the user needs to fill in certain details in the SODAR GUI. SODAR contains multiple built-in templates for, e.g., generic RNA sequencing, germline DNA sequencing and mass spectrometry-based metabolomics. After import or creation, the sample sheets are stored in an object-based format in the SODAR database for easy search and modification. In the GUI, they are presented to the user as spreadsheet-style study and assay tables.

The user can edit sample sheets in the SODAR GUI [Additional File 2]. Cells in the study and assay tables can be edited like in a spreadsheet application. For each column, the project owner or delegate can define the accepted format, value choices, value ranges, regular expressions for accepted values, and other settings depending on the column type. This ensures the validity of data and its compatibility with the study’s requirements and conventions.

SODAR supports ontology term lookup for cell editing. Commonly used ontologies such as Human Phenotype Ontology (HPO) [36], Online Mendelian Inheritance in Man (OMIM) [37], and NCBI Taxonomy Database Ontology (NCBITaxon) [38] can be uploaded into SODAR for local querying as OBO or OWL files, without the need to rely on third party APIs. Manual entering of ontology terms is also allowed. It is possible to include multiple ontology terms in a single cell and one or several ontologies can be used in a single column.

In addition to cell editing, the user can insert and remove rows for study and assay tables. Cells for existing sources, samples, materials, or processes are auto filled by the editor when including a new row. Similarly, if multiple rows contain references to the same entity, all related cells are automatically updated in the tables when modifying them on a single row. SODAR validates all edits using the AltamISA parser [33]. This ensures the validity and ISA-Tab compatibility of the sample sheets at each point of editing.

When editing sample sheets, old sheet versions are stored as backup. These versions can be compared and restored in case of mistakes, as well as exported from the system. SODAR allows for sample sheet export in the full ISA-Tab TSV format, or simplified Excel tables. Replacing existing sheets with versions modified outside of SODAR is also supported.

### 3.4 Integrating SODAR Core based sites

Several subcomponents of the SODAR server such as project and user management have proven to be useful in other contexts. We have extracted them into the SODAR Core software package [32] which forms the foundation of other projects such as VarFish [39] and Kiosc [40]. Using a common library for projects and access management has several advantages and enables the integration of VarFish and Kiosc with SODAR.

SODAR can be configured to work as a “source” site. Applications based on SODAR Core can then be configured as “target” sites of the source site. Projects and access to users will then be synchronized to target sites. This allows us to manage sample and experiment definitions in SODAR and upload corresponding variant data to VarFish. VarFish can then use the REST APIs defined by SODAR for synchronizing sample metadata, such as phenotype terms, directly from SODAR. Similarly, users can upload mass data files into the iRODS data repository and create access tokens to them in SODAR. These tokens can be used to provide data visualization applications in Kiosc with data access via HTTP and iRODS protocols or external applications such as UCSC Genome Browser.

### 3.5 SODAR Administration

We provide a straightforward way to install SODAR and related components (SODAR, iRODS, Davrods, and supporting database servers) and maintain such an installation based on Docker containers and Docker compose. Detailed installation instructions can be found in the “sodar-docker-compose” repository linked to in the section “4.2 Source Code Availability.”

The entire system can be set up using an external LDAP or ActiveDirectory server for users and credentials, or as an alternative in a standalone fashion where SODAR hosts this information. Existing iRODS installations can also be used with SODAR. For administrators, SODAR features dashboards which provide statistics regarding projects and usage of storage resources.

## 4. Data and Source Code Availability

### 4.1 Data Availability

A demonstration instance with data is available at https://sodar-demo.cubi.bihealth.org. All source code is available at https://github.com/bihealth/sodar-server. Example metadata for demonstration projects is available in ISA-tab format at https://github.com/bihealth/sodar-paper.

### 4.2 Source Code Availability

Project name: sodar-server

Project home page: https://github.com/bihealth/sodar-server

Operating system: Linux/Unix

Programming language: Python

License: MIT

RRID: SCR_022175

Biotools: biotools:sodar

## 5. Additional Material

Additional File 1

- File format: Portable Document Format (.pdf)
- Title of data: Data management software comparison table
- Description: Comparison of features between SODAR and related data management software.

Additional File 2

- File format: Portable Document Format (.pdf)
- Title of data: SODAR sample sheet edlevitor
- Description: Figure consisting of screenshots of the SODAR sample sheet editor with its major features annotated.

## 6. Abbreviations

API: Application Programmable Interface
BAM: Binary Alignment Map
BIDS: Brain Imaging Data Structure
CDISC: Clinical Data Interchange Standards Consortium
CUBI: Core Unit Bioinformatics
DMF: Data Management Framework
DRS: Data Repository System
EGA: European Genome-phenome Archive
ELN: Electronic Laboratory Notebook
GEO: Gene Expression Omnibus
GHGA: German Human Genome-phenome Archive
GUI: Graphical User Interface
HDF5: Hierarchical Data Format v5
HPO: Human Phenotype Ontology
HTTP: Hypertext Transfer Protocol
IDS: Instrument-specific Data System
IGV: Integrative Genomics Viewer
IRODS: Integrated Rules-Oriented Data System
ISA: Investigation Study Assay
JSON: JavaScript Object Notation
LDAP: Lightweight Directory Access Protocol
LIMS: Laboratory Information Management System
MIT: Massachusetts Institute of Technology (also commonly used with “MIT license”)
NCBI: National Center for Biotechnology Information
NFDI: Nationale Forschungsdateninfrastruktur (German National Research Data Infrastructure)
OBO: Open Biological and Biomedical Ontologies
OMIM: Online Mendelian Inheritance in Man
OpenBIS: Open Biology Information System
OWL: Web Ontology Language
PAM: Pluggable Authentication Mechanism
PEP: Portable Encapsulated Projects
RBAC: Role-Based Access Control
RDM: Resource Data Management
REST: Representational State Transfer
SDM: Scientific Data Management
SDMS: Scientific Data Management System
SODAR: System for Omics Access and Retrieval
TSV: Tabular Separated Values
UCSC: University of California Santa Cruz
VCF: Variant Call Format
WebDAV: Web-based Distributed Authoring and Versioning)
XML: Extensible Markup Language

## 7. Competing Interests

The authors declare that they have no competing interests.

### 7.1 Funding

This work has been supported by the Ministry of Education and Research (BMBF), as part of the National Research Initiative ‘Mass Spectrometry in Systems Medicine’ (MSCoreSys), grant-ID 031L0220A (MSTARS) and by the Deutsche Forschungsgemeinschaft (DFG, German Research Foundation) – project-ID 427826188 – SFB 1444.

### 7.2 Author’s Contributions

Conceptualization: MN, MH, DB. Funding Acquisition: DB. Methodology: MH, MN. Project Administration: MH, DB. Resources: DB. Software: MN, MH, OS, PP. Supervision: MH, DB. Writing and Editing: all authors

## Acknowledgements

The authors thank all internal users, particularly CUBI members, for their feedback. Some icons from OpenMoji.org were used in the figures.

## Notes

### Competing Interest Statement

The authors have declared no competing interest.

### Summary of Updates

Page 4, 2nd paragraph: typo corrected; Page 7, 2nd paragraph: edited according to reviewer comments; Section 1.5: public repository context clarified; Reference list updated and cleaned up.

