## Additional File 1 (Supplemental Table) for "SODAR: managing multi-omics study data and metadata"

| (yes)=can be implemented within framework | | SDMS | | | | ELN | DRS | | DMF | |
|---|---|---|---|---|---|---|---|---|---|---|
| | | SODAR | qPortal | FAIRDom Seek | OpenBIS ELN-LIMS | eLabFTW | Dataverse | Yoda | Molgenis | Zendro |
| **SDMS features** | | | | | | | | | | |
| 1. Overarching | | | | | | | | | | |
| 1.a | Structure into projects/folders | yes | yes | yes | yes | yes | yes | yes | yes | yes |
| 1.b | Access control | yes | yes | yes | yes | yes | yes | yes | yes | yes |
| 1.c | Automation possible via API | yes | yes | yes | yes | yes | yes | yes | yes | yes |
| 1.d | Open standards / formats | yes | no | yes | no | yes | yes | yes | yes | yes |
| 2. Planning | | | | | | | | | | |
| 2.a | Structured recording of assays/experiments | yes | yes | yes | yes | limited | no | no | (yes) | (yes) |
| 2.b | Flexible definition of studies/experiments | yes | limited | yes | limited | yes | no | no | (yes) | (yes) |
| 2.c | Controlled vocabulary | yes | yes | yes | yes | no | no | no | (yes) | (yes) |
| 2.d | Ontologies | yes | no | yes | no | no | no | no | (yes) | (yes) |
| 3. Data collection | | | | | | | | | | |
| 3.a | Storage of files possible | yes | yes | yes | yes | yes | yes | yes | (yes) | no |
| 3.b | Many / large files | yes | no | no | limited | no | yes | yes | (yes) | no |
| 4. Data analysis | | | | | | | | | | |
| 4.a | Meta data API | yes | no | yes | yes | yes | yes | yes | yes | yes |
| 4.b | Mass data files API | yes | no | yes | limited | no | yes | yes | no | no |
| **5. Further features** | | | | | | | | | | |
| **5.a ELN** | | | | | | | | | | |
| 5.a.i | Flexible data entry text/table/pictures | no | no | no | yes | yes | no | no | no | no |
| **5.b DRS** | | | | | | | | | | |
| 5.b.i | Host public data repositories | no | no | yes | no | no | yes | yes | (yes) | (yes) |
| **5.c DMF** | | | | | | | | | | |
| 5.c.i | Easy creation of tables | no | no | no | no | no | no | no | yes | yes |
| 5.c.ii | User-centric data entry masks | limited | yes | limited | limited | limited | no | no | yes | yes |
| 5.c.iii | Predefined components, e.g., for data analysis | no | no | no | yes | no | no | no | yes | no |
