## Supplementary figures and images for "SODAR: managing multi-omics study data and metadata"

### Additional File 2 (Supplemental Figure)

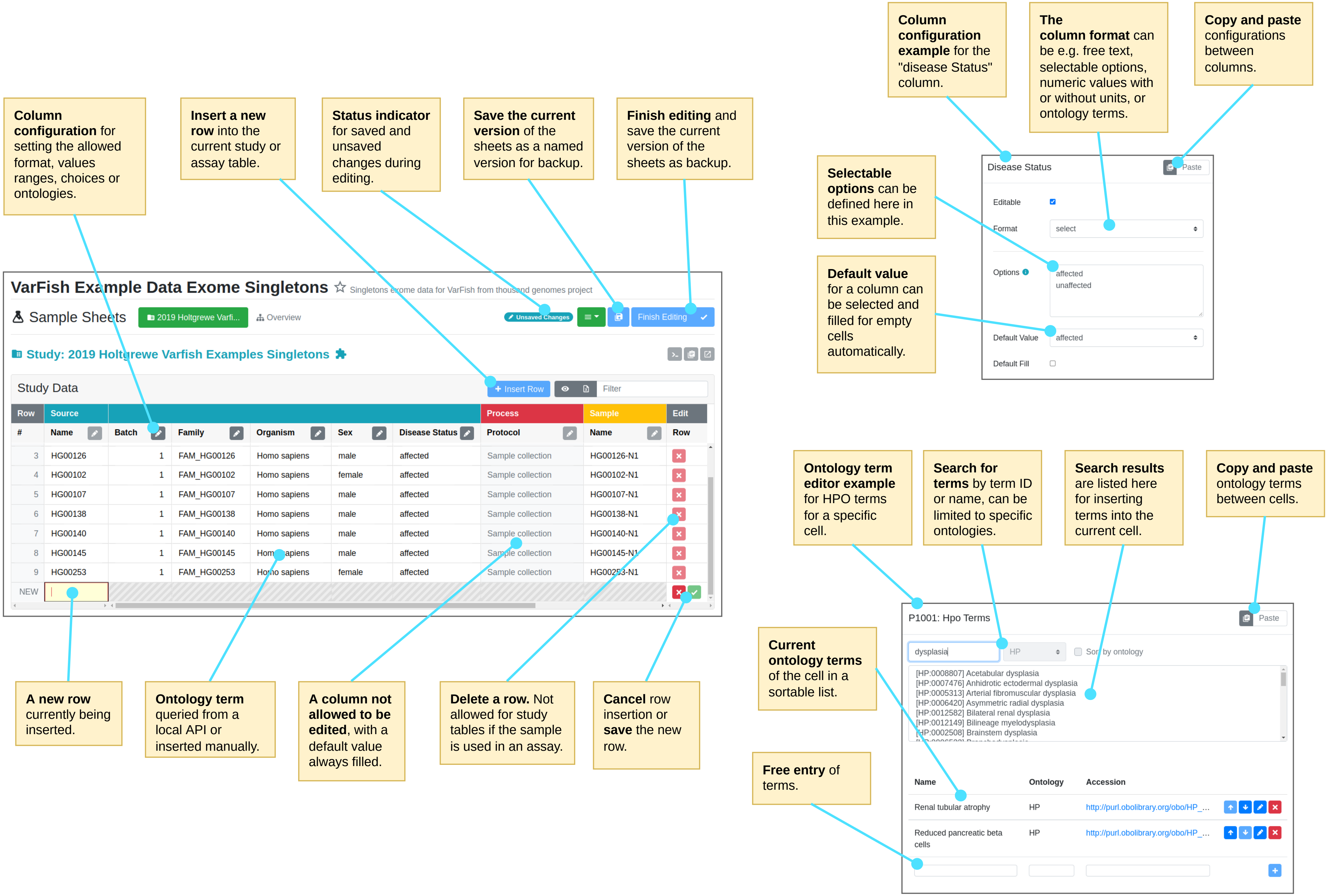
